# A psychometric platform to collect somatosensory sensations for neuroprosthetic use

**DOI:** 10.1101/2020.07.23.218222

**Authors:** Giacomo Valle, Francesco Iberite, Ivo Strauss, Edoardo D’Anna, Giuseppe Granata, Thomas Stieglitz, Stanisa Raspopovic, Francesco M. Petrini, Paolo M. Rossini, Silvestro Micera

## Abstract

Somatosensory neuroprostheses exploit invasive and non-invasive feedback technologies to restore sensorimotor functions lost to disease or trauma. These devices use electrical stimulation to communicate sensory information to the brain. A sensation characterization procedure is thus necessary to determine the appropriate stimulation parameters and to establish a clear personalized map of the sensations that can be restored. Several questionnaires have been described in the literature to collect the quality, type, location and intensity of the evoked sensations, but there is still no standard psychometric platform. Here we propose a new psychometric system containing previously validated questionnaires on evoked sensations, which can be applied to any kind of somatosensory neuroprosthesis. The platform collects stimulation parameters used to elicit sensations; records subjects’ percepts in terms of sensation location, type, quality, perceptual threshold, and intensity. It further collects data using standardized assessment questionnaires and scales, performs measurements over time, and collects phantom limb pain syndrome data. The psychometric platform is user-friendly and provides clinicians with all the information needed to assess the sensory feedback. The psychometric platform was validated with three trans-radial amputees. They platform was used to assess intraneural sensory feedback provided through implanted peripheral nerve interfaces. The proposed platform could act as a new standardized assessment toolbox to homogenize the reporting of results obtained with different technologies in the field of somatosensory neuroprosthetics.

## Introduction

Somatosensory neuroprostheses are highly innovative devices [1]. Several research groups have investigated the ability to restore sensory feedback in patients with upper or lower limb amputation, tetraplegia or paraplegia using invasive [2–11] and non-invasive [12–15] interfaces with the PNS and the CNS (Figure 1). The main aim of these technologies is to elicit somatotopic referred sensations emanating from the affected limb, creating a personalized map of the these sensations which could be used as sensory feedback aimed at improving the patients’ quality of life[16,17]. All these approaches use neural stimulation to evoke sensations stemming from contact with sensory peripheral nerves or the neural interfaces are placed directly on the somatosensory cortex.

**Figure 1:**
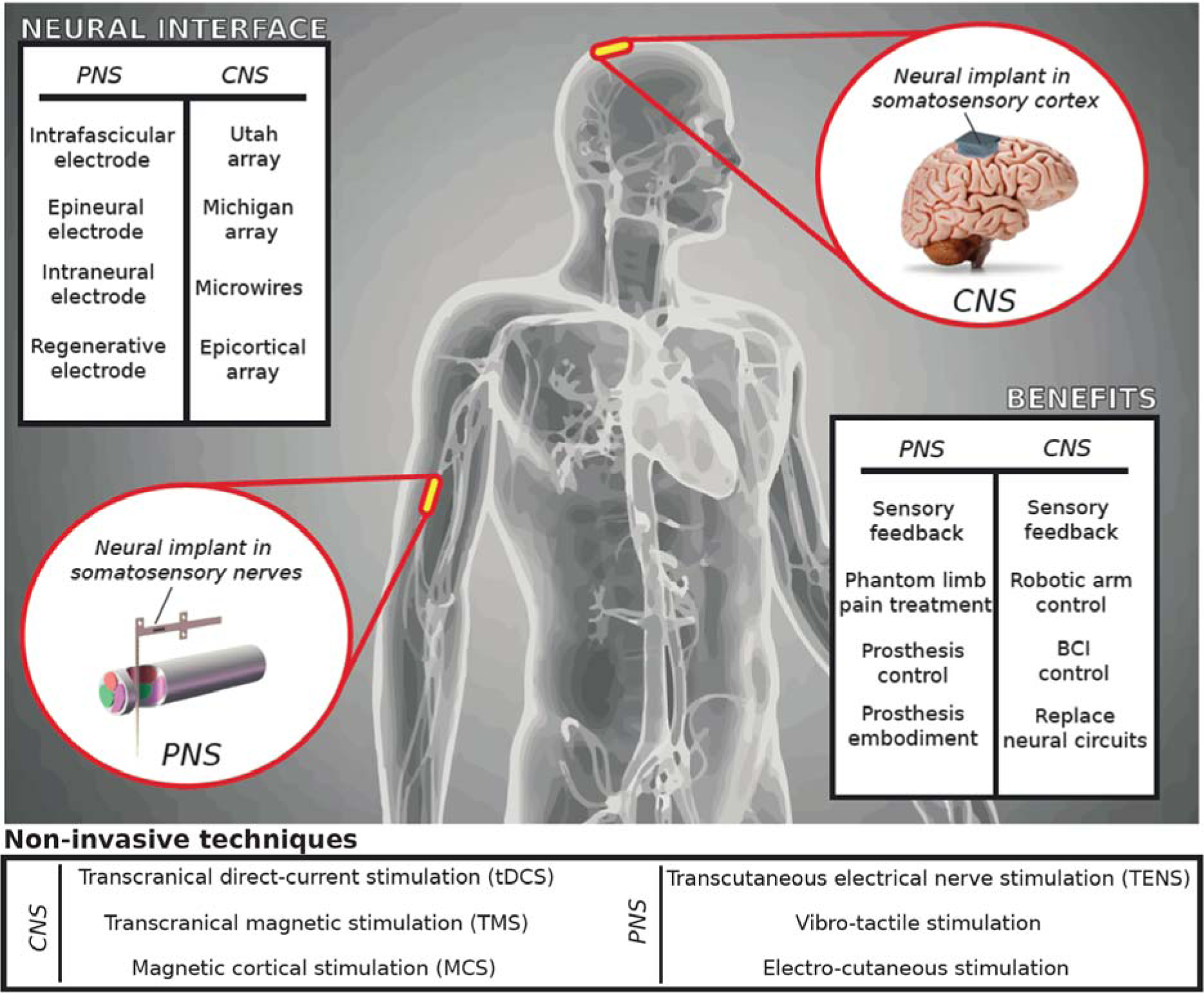
Neuroprosthetic applications. Neurotechnologies for restoring somatosensations have been developed for peripheral (PNS) or central (CNS) nervous systems. The stimulation technique used to restore sensory feedback can be invasive (surgically implanted and in intimate contact with the nervous tissue) or non-invasive (applied on the skin surface). Delivering a stimulation to the brain or peripheral nerves provides benefits such as the control of robotics, smart prosthetics, or other assistive technologies.

Since there is inter-subject variability due to the different nerve structures, implantation levels and innervation[18], together with the subjective perception of the elicited sensations, a “sensation characterization” procedure is necessary to obtain a uniform sensation mapping (Figure 2). The goal of this procedure is to collect all the stimulation parameters corresponding to the evoked sensations characterized by the intensity, quality, location and type in order to have a clear sensation map. The mapping phase is crucial to implement an effective real-time assistive system, e.g. bidirectional hand or leg prostheses, eliciting homologous referred sensations emanating from the phantom limb (somatotopic) for therapeutic or functional purposes. In fact, the personalized sensation map is often translated into a robotic arm or hand in order to elicit sensations during object manipulation tasks aimed at increasing patient motor control performance[2,4,19,20]. When the patient is controlling a robotic arm, and touches a surface with the second robotic digit, the sensation perceived should be in the same location (index), with the safety and exact intensity (mapped with the pressure force of the robotic finger) and the type should be line with finger pressure (i.e. no electricity or warmness). The personalized sensation map should thus be as detailed as possible.

**Figure 2:**
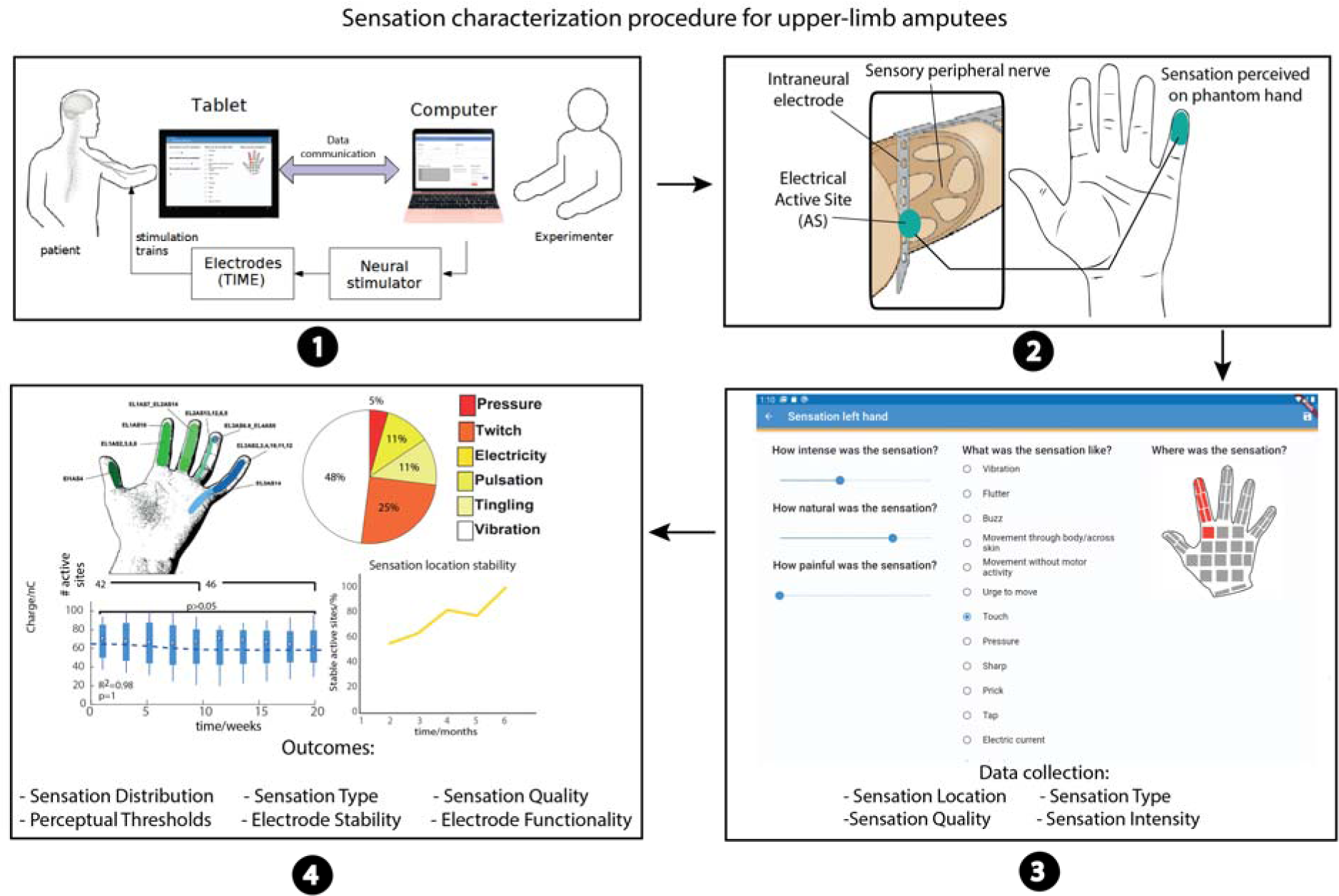
Sensation characterization procedure. (1) Stimulation parameters are selected. The stimulation trains are delivered using the neurostimulator, the software also sends control commands to the Easy Quest app. (2) Patient perceives a stimulation-evoked sensation on the phantom hand thanks to the neural implant. (3) Easy Quest app in ODF mode is used to report the sensations (4) Experimenters collect all sensation characterization outcomes and import them in Matlab or Excel to plot the results.

Several psychometric questionnaires exist regarding the quality and type of the sensations evoked[15,21–23]. However, they do not appear to be easy-to-use or fast for recording and integrating all the properties of the elicited sensations with detailed standard questionnaires, and which could be used for several types of sensory feedback.

The psychometric platform presented in this study provides a uniform way of characterizing and quantifying the artificial sensory feedback systems used for invasive and non-invasive, peripheral and central sensory feedback, in order to efficiently compare, optimize and evaluate all the different approaches. Our platform records the stimulation parameters, quality, type, intensity and location of the evoked sensations. All the sensation data are collected from questionnaires already presented in the literature.

The platform also provides a user-friendly graphical user interface with a touch screen for the patient’s answers that not only enables the patient to describe the percept in detail, but also provides clinicians with all the main information on the evoked sensation. The platform accepts new questionnaire definitions as text, and is easy to understand and implement. This means that researchers can add new questionnaires, such as phantom limb pain (PLP)[21,22], in order to collect information on new treatments.

This psychometric platform was tested on three trans-radial amputees who had an intrafascicular electrode [24] implanted in their median and ulnar nerves for six months each. The patients responded using the psychometric platform when they received electrical stimulation by the electrical contacts of the neural interfaces. The software was used by clinicians and engineers to collect the data. This has proven to be more convenient than writing down all the answers in weekly trials over 18 months.

In this study, we describe the usability of this new platform. We believe that our new psychometric platform will facilitate and unify the characterization of percepts and the comparison of the effects when applying different neural stimulation techniques or using different devices.

## Methods

### Software platform

The psychometric platform is made up of a mobile application for compiling questionnaires (which we have called Easy Quest), two desktop tools (Easy Quest Create and Easy Quest Evaluate) and a desktop application to control the neurostimulator and, also a mobile app.

The Easy Quest mobile app is described in depth in the following sections.

Easy Quest Create shows a simple graphical user interface in which the experimenter can create a list of questions from a set of pre-defined types. The content can be customized. Easy Quest Evaluate is devised for the rapid evaluation of a set of answers, the software reads the archive file exported by Easy Quest and exports a CSV file. The choice of CSV format of the results makes further analyses easier, as it is compatible with Matlab (The MathWorks, Inc., Natick, Massachusetts, United States), and Microsoft Excel (Microsoft Corporation, Redmond, Washington, United States).

The desktop application for the actual neurostimulation is not described here, because its design is strongly dependent on the type of experiment and neurostimulation device (communication protocols, stimulator commands and architecture), however it is mentioned as part of the experimental setup.

### Somatosensory questionnaires

Somatosensory descriptors were selected from the literature and clinical settings also including questionnaires that have already been used in neuroprosthetic studies. Several options describing the type, quality, intensity and the location are presented in order to characterize the somatosensory percepts being evoked during the stimulation. To describe the quality of sensations, we used a scale presented by Lenz et al.[22] and used also by Valle et al.[9]. For the sensation type, we adapted the questionnaire proposed by Kim et al.[21] based on our experience with several upper limb patients stimulated with invasive[2,9,25–31] and non-invasive technologies[12,32]. We also considered other studies on sensations elicited using peripheral[3,29,33] or central[4] neural stimulation approaches. For the intensity, we used a Visual Analogue Scale (VAS)[34] already presented by Tan et al.[35]. Lastly, the perceived sensation locations were shown directly on a schematic representation of the human hand. it is further possible to select the feet, arms or legs [10,11] with several possible spots (Figure 2). In this way, the patient can accurately indicate the affected areas.

We added several questionnaires in order to collect information on phantom limb pain (PLP): visual analogue scale (VAS)[34] and neuropathic pain symptom inventory (NPSI)[36]. It is also possible to add or modify the existing questionnaires in order to adapt the platform to the needs and specifications of the clinical trial.

### Use cases

Two main use cases for the app were identified (Figure 3). In the first, the user fills in a questionnaire and saves the results on the device, defined as the “local fill-in” (LF). In the second, an external software prompts the app to show a questionnaire and to send back the results, defined as “on demand fill-in” (ODF). The two cases (Figures 3A, B) involve the same procedure in the part where the user is asked to fill in the answers.

**Figure 3:**
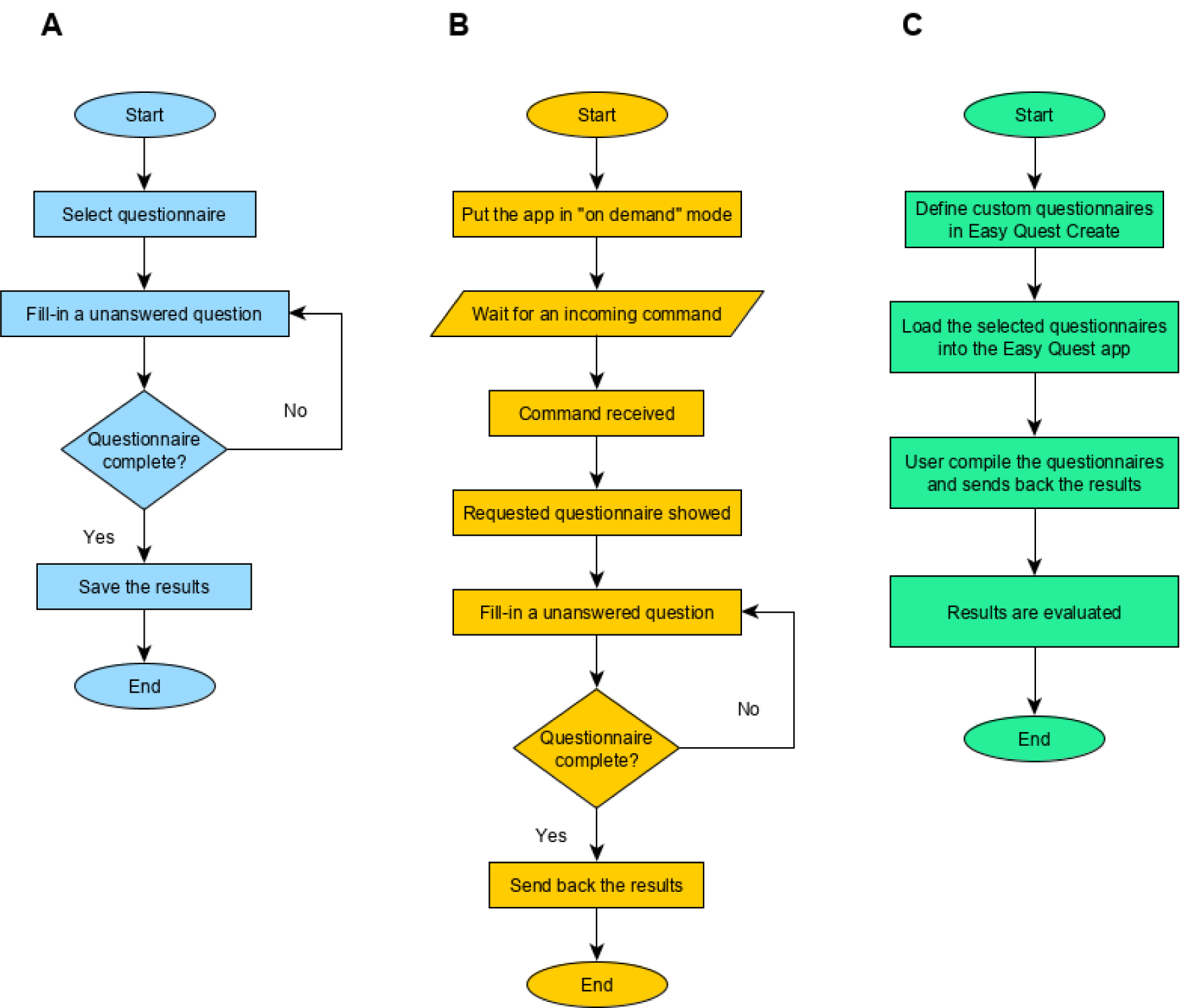
Use cases. The three main features of the psychometric platform, the first two are implemented by the mobile app, the last by the whole system. **A** Defined as Local Fill-in (LF), where the users compile a questionnaire and the answers are stored in the device. **B** On demand fill-in (ODF), in this case the app waits for an external command from a controller app containing information on the questionnaire to be shown; the fill-in procedure is the same but nothing is stored within the device, instead results are sent back to the controller. **C** The procedure seen from the experimenter’s point of view, here the role of the other software programs of the platform (Easy Quest Create, Easy Quest Evaluate) is explained.

The main difference, besides the location where the results are stored, is how the procedure starts: in the first case, the user choses a questionnaire by selecting it from the main menu, in the second, the app waits for an external command, usually from the network, instructing the software to show a specific questionnaire.

The application can set recurrent reminders for specific questionnaires, enabling the experimenter to plan the follow-up for home use by the patient, and these reminders prompt the user to fill-in the questions in LF mode.

A third use case (Figure 3C) explains the workflow from the perspective of the experimenter, who uses the companion software to define new questionnaires at the beginning of the experimentation and to display the results at the end.

### Software architecture

The software was developed in Dart, an object-oriented programming language developed by Google in order to address server-side, web and mobile platforms. The mobile SDK, Flutter, compiles the code in fast native apps for Android and iOS devices.

The app is developed following the MVC (Model View Controller) pattern, and a simple ORM (Object-Relational Mapping) is implemented to store the models in an SQLite database in the device’s memory. The ORM is accessed through classes which show APIs where serialized objects can be stored and retrieved.

To implement the ODF, a simple HTTP server runs in background thus the app can, when requested, wait for remote commands from the network. While doing so, the app shows a numerical code, which must be notified to the experimenter to secure the remote connection.

An interface with the mail app is used to send the completed questionnaires as a CSV file by email.

Another provider class parses the questionnaires defined in JSON (JavaScript Object Notation) format, making it possible to create and add new questionnaires to working devices, without code interventions and recompiling the whole app. The import service can parse a compressed file containing a set of questionnaires and also a collection of images referred to in the questions. There are five questions accepted by the parser: (1) open, which prompts the user for a string (2); radio, which asks the user to choose one option from a set (3); multiple-choice; (4) slider, where users have to select a number or a label with a slider (5) image touch, where the user selects a set of touchable areas displayed on top of a given background image.

The app enables multiple users to access the same device while keeping the results separate.

The system architecture is shown in Figure 4, along with the external software highlighting its relationship with the app modules.

**Figure 4:**
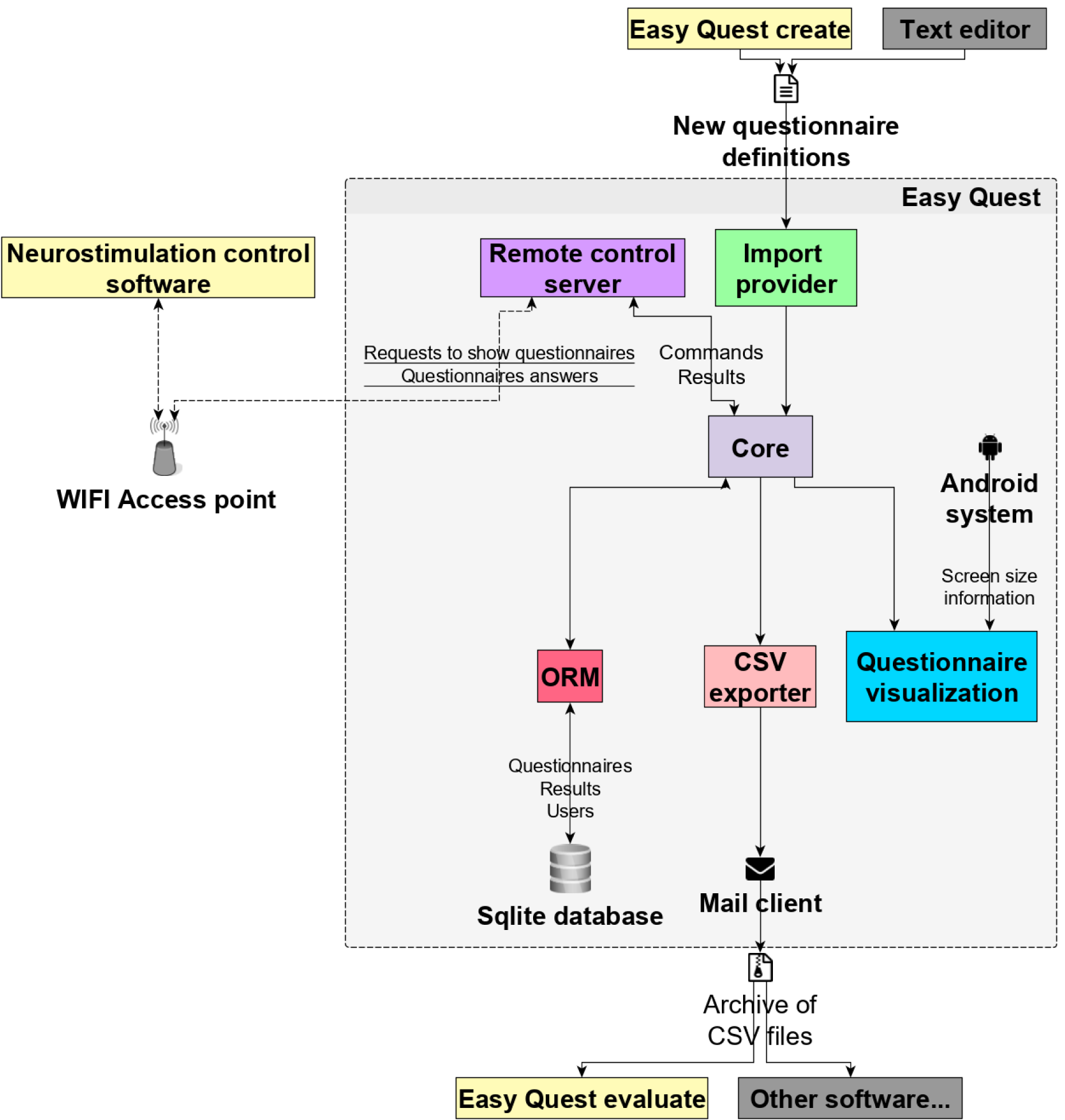
Software architecture. The main components of the platform depicted as squares, external services are shown with an icon and communication with arrows, some show a label with examples of the information flowing through. A grey shadow surrounds the software modules of the mobile app (Easy Quest).

The app UI/UX is designed in accordance with Material, an open source system of guidelines developed by Google. The view layer written for the app exploits all the available space, presenting the questionnaire as a list of questions on small devices and as a grid on larger screens.

### Quality and usability assessment

During the clinical trial we collected feedback information from patients, clinicians and engineers who used the platform presented in this study in three clinical trials (N=12). The investigations regarded the development and assessment of bidirectional hand prostheses for upper limb amputees with a neural sensory feedback delivered by implantable electrodes[9,26–28,31]. After six months of use, we asked participants to answer different quality and usability questions using: questionnaires for user interface satisfaction (QUIS)[37], system usability scales (SUS)[38,39], Nielsen’s attributes of usability (NAU)[40] and after-scenario questionnaires (ASQ)[41]. We collected and analyzed all the information using validated and standardized questionnaires (Figure 5).

**Figure 5:**
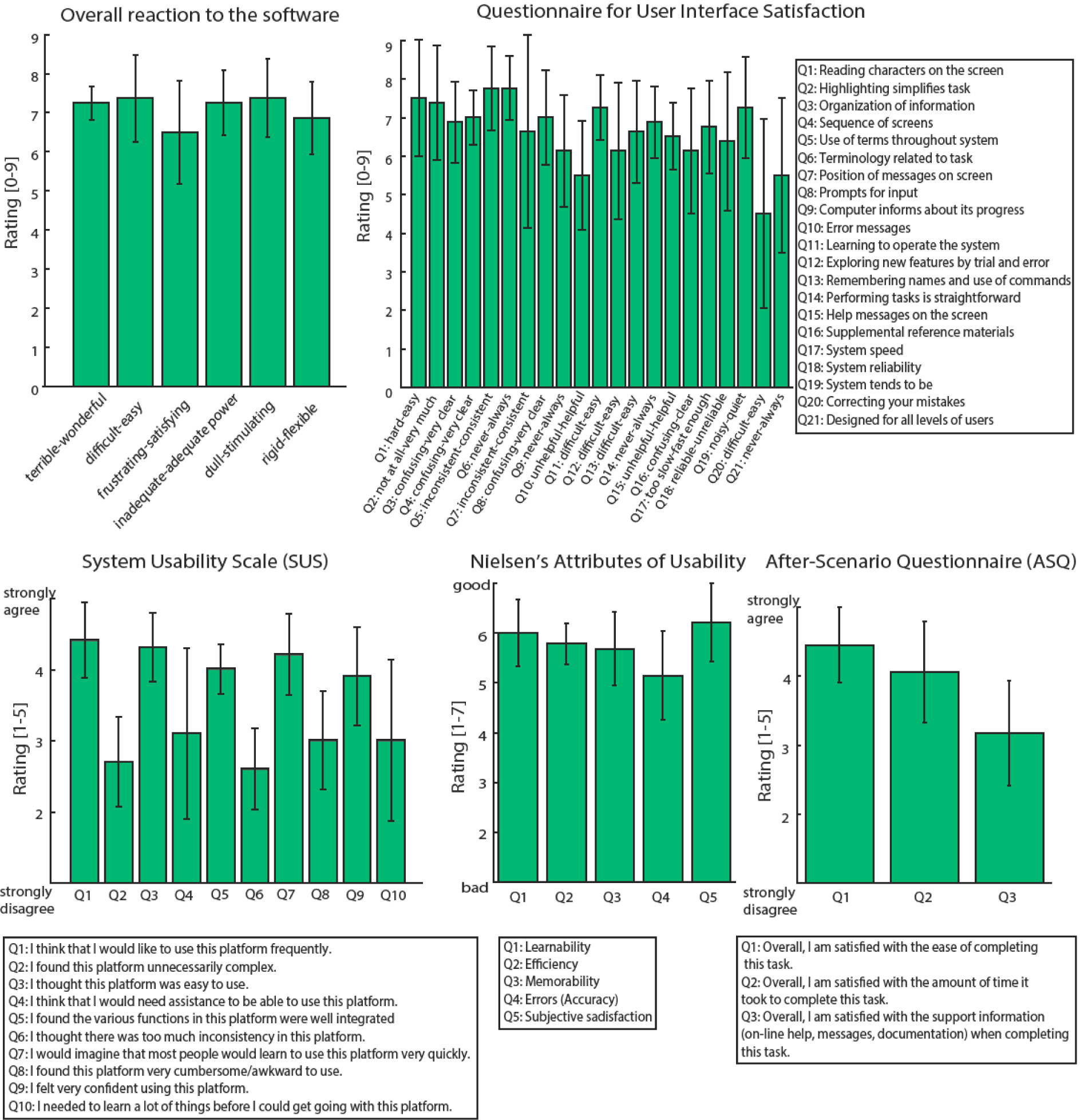
Usability assessment. All the usability scales are reported: Overall reaction to the software, QUIS, SUS, NAU and ASQ. Three clinicians, six engineers and three patients evaluated the psychophysical platform (N=12). The data in the figure are represented as means ± standard deviations.

## Results

### Somatosensory questionnaires for sensation characterization

To efficiently characterize the sensations emanating from (invasive or non-invasive) electrical (central or peripheral) stimulation, a user-friendly platform is needed with a set of somatosensory related questionnaires. This helps to reduce the long time required to collect all the electrically-evoked sensation data.

To assess the properties of the sensations being evoked by stimulating peripheral nerves using a neural interface in trans-radial amputees, we used the psychometric platform presented here. We performed a procedure called “sensation characterization” with all the patients involved in the clinical investigation (Figure 2). For each electrically active site used to stimulate the nerve, the neural stimulation was delivered, and the patient was asked to report the sensations he/she felt. This mapping phase enabled us to identify the sensation properties for all the stimulation channels of the implanted electrodes by varying the stimulation parameters and building a personalized map of the sensations. The stimulation parameters varied in terms of frequency (1-1000 Hz), pulse-width (1-120 _μ_s), and amplitude (1-1000 _μ_A), as well as stimulation train duration (discrete or continuous). We collected the sensation intensity, quality, type and location of the patient’s perceived sensations.

The intensity was used to find the perceptual thresholds for each stimulation channel[2,28,31], together with the range of stimulation (between threshold and below pain level). Using a VAS scale in the range from 0 to 10 also enables us to identify perceptual magnitude levels[3,31,42].

The quality of the sensory feedback was assessed in order to test different stimulation strategies and approaches[9,31], since this quality is considered to be an important factor for prosthesis acceptance[43]. To quantify the perception quality and naturalness, we used a scale[22] from 1 (totally unnatural) to 5 (totally natural).

The type of sensation was collected in order to understand the type of fibers being recruited during the stimulation and to identify the best channels for restoring homologous sensations while using the bidirectional prosthesis. We used 20 descriptors (Table 1) considering all the important aspects. In this platform the patient could also report a new sensation or add comments in an empty text box when a correct descriptor for the elicited sensation was lacking.

**Table 1:** List of sensory descriptors. Here the chosen descriptors are shown, the user can also enter free text when the other options are insufficient.

|  |
| --- |
| Vibration |
| Flutter |
| Buzz |
| Movement through body/across skin |
| Movement without motor activity |
| Urge to move |
| Touch |
| Pressure |
| Sharp |
| Prick |
| Tap |
| Electric current |
| Shock |
| Pulsing |
| Tickle |
| Itch |
| Tingle |
| Numb |
| Warm |
| Cool |

The sensation location was reported using a picture of the limb of interest (foot, arm, leg or hand) with several highlighted spots (20 for foot, 24 for leg, 48 for arm and 45 for hand) (Figure 2). The zones with a higher density of receptors had more selectable spots. This information is useful to understand the electrode stimulation selectivity (analyzing the spreading of the zone) and the layout of the fibers inside the nerve. In addition while the bidirectional prosthesis was being used, the location map was needed to stimulate the correct active sites eliciting the somatotopic sensation during the prosthesis hand/finger contact with objects[2].

Finally, several questions can be used to assess phantom limb pain levels before and after a pain treatment with electrical stimulation[28]. We decided to use two different questionnaires (VAS and NPSI) to characterize the location, quality and intensity of the pain[10,28].

### Software usability

The usability testing of the app was performed on an Android phone (a Nexus 6p), designed by Huawei and running Android 8.

The app loading time is less than two seconds, needing only the time to open the local database, and after the login screen, the user can access all the main functions in no more than two taps.

The home page shows a list of all the available questionnaires, the user can tap on each one to see the questions and fill in the answers, which are stored in the internal database. From the lateral menu (drawer), the ODF mode can be accessed in only one tap, after which the app will wait for a network command containing the identifier of the questionnaire to be shown.

Minimal user interaction is needed to complete a questionnaire, usually all the questions need just one tap, except for the multiple choice and clickable area ones. The average time to fill-in a sensation characterization questionnaire is 10 seconds.

The export page lets the user write all the stored data in a CSV archive file and opens the default mail to send to the experimenters for further analysis, facilitating and speeding up the data gathering phase.

A specific section of the app lets the user choose which questionnaire should be visible in the home page, personalizing the user interface for a specific use.

Other pages are designed for secondary tasks, such as previewing stored answers and editing settings.

### Psychometric system validation

In order to assess the usability and quality of this novel psychometric platform to collect somatosensory percepts, several questionnaires were filled in by different kinds of users. Three patients, six engineers and three clinicians evaluated the system by answering four questionnaires after using the platform in clinical applications (Figure 5). Analyzing the results, the overall reactions to the system were very positive. The average score was 7.1± 0.3. Considering the user interface satisfaction (QUIS), the rating achieved was 6.6±0.8. In both these questionnaires the maximum achievable score was 9.

In the SUS (range 1-5), Q1-Q3-Q5-Q7-Q9 scored 4±0.2, while Q2-Q4-Q6-Q8 scored 2.3±0.2. These results indicate that the users agreed more with the positive sentences and disagreed more with negative ones. The NAU (range 1-7) showed high ratings of 5.6±0.5, and the ASQ (range 1-5) showed an average value of 3.7±0.8.

During the clinical trial, the psychophysical platform was used over 1000 times.

## Discussion

Electrical stimulation has been proposed as a way of restoring somatosensations[15,43– 46] in cases where they have been lost due to injury or disease in both the CNS or the PNS. In fact, sensory feedback is crucial to improve the motor control of robotic limbs or prostheses, enabling the patient to be more efficient in manipulating objects[2,19,47]. The sensations evoked thus had to be characterized in detail in patients receiving stimulation in order to restore the sensory information. The psychometric questionnaires were able to register all the aspects of the sensations being restored in a reliable and efficacy way, considering more descriptors than in previous studies[22] and using a user-friendly platform.

Currently, there are many important sensation properties which need to be collected in order to obtain an intuitive and rich sensory feedback. In particular, the sensation location, type, quality and intensity are valid and extendable for all the approaches in different neurological conditions. Considering the previously presented interface to collect stimulation-evoked somatosensory percepts, Geng and collaborators[23] showed a platform used to evaluate electrical stimulation to relieve Phantom Limb Pain. Their platform was interfaceable with one type of neural stimulator and contained three questions to characterize the evoked sensation considering 12 sensation descriptors. The psychometric platform presented here reports somatosensory percepts based on five questionnaires containing 20 standard sensory descriptors (Table 1). The platform exploits a customizable, fast and easy to use GUI which can be efficiently connected to several neural stimulators[28,48,49].

Since several groups are currently using electrical stimulation to restore sensory feedback, a standard somatosensory platform could facilitate their comparison, assessment and optimization. Our findings support the conclusion that this psychometric platform could help and accelerate the development of sensorimotor neuroprostheses.

Given the simple software architecture, this platform is flexible in terms of modifications and upgrades. It is possible to add new questionnaires regarding other aspects of sensory feedback restoration. For example, two important features to be considered for the development of the next generation of somatosensory neuroprostheses are embodiment[26,50] and psychological/affective aspects[51].

The psychometric platform is simple to interface with other devices and also with existing software, thanks to its open and platform-agnostic interfaces: in ODF mode the HTTP interface accepts commands regardless of the device and the programming language of the sender application (all major languages can implement HTTP communication effortlessly). Answers to the questionnaires are exported in a CSV format, making it easy for any other software program to import and analyze them.

Considering the results of the usability assessments (Figure 5), users highlighted various positive and negative aspects which will then help us to improve the platform. The most positive aspect in terms of the ‘overall reaction to the software’ was that the software is easy to use, which is crucial both for patients and experimenters.

The QUIS answers revealed that this system is consistent and very clear, however we still need to improve error and warning messages. These aspects mainly regard the experimenters’ side. The SUS again indicated that the system is easy to use and intuitive, but additional material and instructions should be included as support. Also the NAU showed a high user satisfaction along with a request for more error messages. Finally, the ASQ revealed ‘the ease of completing this task’, thus highlighting the need for more support, information and documentation. We thus intend to improve the platform using these usability results.

### Study Limitations

There are several limitations connected to the patient attention at the time of testing. To solve this issue, it is important to repeat the test multiple times over multiple days in order to increase its reliability. The test is also highly subjective, and the mapping results could strongly depend on the sensation of the patient and his / her personal experience[52]. The individual subjective differences remain a big challenge for interpreting the somatosensory results and also the semantic differences.

Sham (placebo) and blind stimulations could also be delivered to test individual response bias and identify possible unreliable self-reports.

The software design, particularly the GUI, was inspired by the principles of the ISO 9241 standard. In fact, the users’ opinions of the platform were taken into consideration during the design phase and the assessment.

The software will be actively used during experiments and the user experience will be monitored to improve new versions, ensuring an iterative development driven by user feedback, as also stated in ISO 9241.

### Conclusions

This study has presented a psychometric platform used to record a complete somatosensory percept description, which can be evoked by several different methods of electrical stimulation in humans. The subjective somatosensory sensation type, location, quality and intensity are collected and used to develop a somatosensory questionnaire, which can be used for neuroprosthesis calibration and optimization. The psychometric toolbox is implemented in a user-friendly software program. The platform was validated in patients with electrodes implanted in the PNS.

We believe that this new somatosensory psychometric system will help to establish a standard and uniform methodology of subjective sensory reports, which is a pivotal step to uniformly develop, adapt and improve somatosensory neuroprostheses.

## Acknowledgments

The funder had no role in the experimental design, analysis, or manuscript preparation or submission. The funder provided funds to complete the study, including investigator salaries, equipment costs, and research and clinical costs. All authors had complete access to data. All authors authorized submission of the manuscript, but the final submission decision was made by the corresponding authors.

## Author contributions

G.V. designed the study, developed the software, analyzed the data, and wrote the paper; F.I. developed the software, analyzed the data and reviewed the manuscript; I.S. designed the study, developed the software, and reviewed the manuscript; E.D. developed the software, and reviewed the manuscript; G.G. and P.M.R. tested the platform with patients; T.S developed the TIME electrodes; T.S. and P.M.R. discussed the results and reviewed the manuscript; S.R. designed the platform; F.P. designed the platform and discussed the results; S.M. designed the study, discussed the results and reviewed the manuscript. All the authors read, commented, and approved the manuscript.

## Data availability

the datasets generated during and/or analyzed during the current study are available from the corresponding author on reasonable request. The Easy Quest Android application is available from the Mendeley Data platform: https://data.mendeley.com/datasets/m85f5dxw4f/draft?a=5ab32d12-6d46-4465-ac9a-43a3813f3f82

## Ethical approval

Ethical approval was obtained by the Institutional Ethics Committees of Policlinic A. Gemelli at the Catholic University, where the surgery was performed. The protocol was also approved by the Italian Ministry of Health. Informed consent was signed. The clinical trial’s registration number on the online platform (www.clinicaltrials.gov) is NCT02848846.

## List of abbreviations

App: Application
ASQ: After-Scenario Questionnaire
CNS: Central Nervous System
CSV: Comma-Separated Values
FDA: Food and Drug Administration
GUI: Graphical User Interface
HTTP: Hyper-Text Transfer Protocol
JSON: JavaScript Object Notation
LF: Local Fill-in
MDR: European Medical Device Regulation (MDR)
MVC: Model View Controller
NAU: Nielsen’s Attributes of Usability
NPSI: Neuropathic Pain Symptom Inventory
ODF: On Demand Fill-in
ORM: Object-Relational Mapping
PLP: Phantom Limb Pain
PNS: Peripheral Nervous System
Q-(1-9): Question-number (#)
QUIS: Questionnaire User Interface Satisfaction
SUS: System Usability Scale
UI/UX: User Interface/User Experience
VAS: Visual Analog Scale

